# DeGenPrime provides robust primer design and optimization unlocking the biosphere

**DOI:** 10.1101/2023.08.11.553048

**Authors:** Bryan Fulghum, Sophie Tanker, Richard Allen White

**Author notes:** To whom correspondence should be addressed: Richard Allen White III, UNC Charlotte.

## Abstract

Polymerase chain reaction (PCR) is the world’s most important molecular diagnostic with applications ranging from medicine to ecology. PCR can fail because of poor primer design. The nearest-neighbor thermodynamic properties, picking conserved regions, and filtration via penalty of oligonucleotides form the basis for good primer design.

DeGenPrime is a console-based high quality PCR primer design tool that can utilize MSA formats and degenerate bases expanding target range for a single primer set. Our software utilizes thermodynamic properties, filtration metrics, penalty scoring, and conserved region finding of any proposed primer. It has degeneracy, repeated k-mers, relative GC content, and temperature range filters. Minimal penalty scoring is included according to secondary structure self-dimerization metrics, GC clamping, tri- and tetra-loop hairpins and internal repetition.

We compared PrimerDesign-M, DegePrime, ConsensusPrimer, and DeGenPrime on acceptable primer yield. PrimerDesign-M, DegePrime, and ConsensusPrimer provided 0%, 11%, and 17% yield respectively for alternative iron nitrogenase (*anfD*) gene target. DeGenPrime successfully identified quality primers within the conserved regions of the T4-like phage major capsid protein (*g23*), conserved regions of molybdenum-based nitrogenase (*nif*), and its alternatives vanadium (*vnf*) and iron (*anf*) nitrogenase. DeGenPrime provides a universal and scalable primer design tool for the entire tree of life.

## 1. Introduction

Polymerase-chain reaction (PCR) is the most important fundamental tool for molecular diagnostics, genetic analysis, viral load testing, molecular biology, phylogenetics, and a plethora of other disciplines. A major cause of failure in PCR experiments is a poor choice in primers. Rules of good design for PCR primers are well-established; they should be 15-30 base pairs (bp) long, without complementary ends, contain between 40-60% guanine (G) or cytosine (C), have minimal dinucleotide repeats, and similar melting temperatures [1, 2]. Primer3 is the gold-standard tool for finding excellent candidate primers for single gene sequences [3, 4]. This tool was not designed to find primers for the multi-sequence alignments (MSAs) often used in phylogenetic studies and does not support the addition of degenerate bases.

When comparing closely related species, there will be some conserved regions of DNA [5]. In fact, as two species are more distantly related, the conserved regions tend to disappear [6]. Conserved regions are the optimal locations for PCR experiments as there can be no primer matching bias or mismatches when the target region is identical across all sequences [2].

DeGenPrime aims to find primers for MSAs based on the conserved regions found across the sequences without relying on any reference, instead using the general principles of good primer design. Beyond a conserved region approach, DeGenPrime also utilizes a filtration digital module that hard filters primers based on limited degeneracy. Our software can build primers independently without bias in a highly scalable manner resolving the diverse and continuously evolving biosphere.

## 2. Program Design

DeGenPrime provides an overall tool kit for primer design via filtration or conserved region picking (**Fig 1**). DeGenPrime input formats include a nucleotide fasta file or a nucleotide MSA with preprocessor tags in clustal format. User tags are processed and stored as static global variables for access throughout various parts of the program. After loading input data, the program checks the sequence file for selected format either a collection of sequences that aren’t aligned, a previously aligned collection of sequences, or a single gene sequence. The quality controls list whether the sequence file is correctly formatted, aligned, and/or an empty file. If it is not aligned, the program aligns the sequences via MAFFT (Multiple Alignment fast Fourier transform) either using global or local based on user specifications with a maximum iteration of 1000 [7].

**Figure 1:**
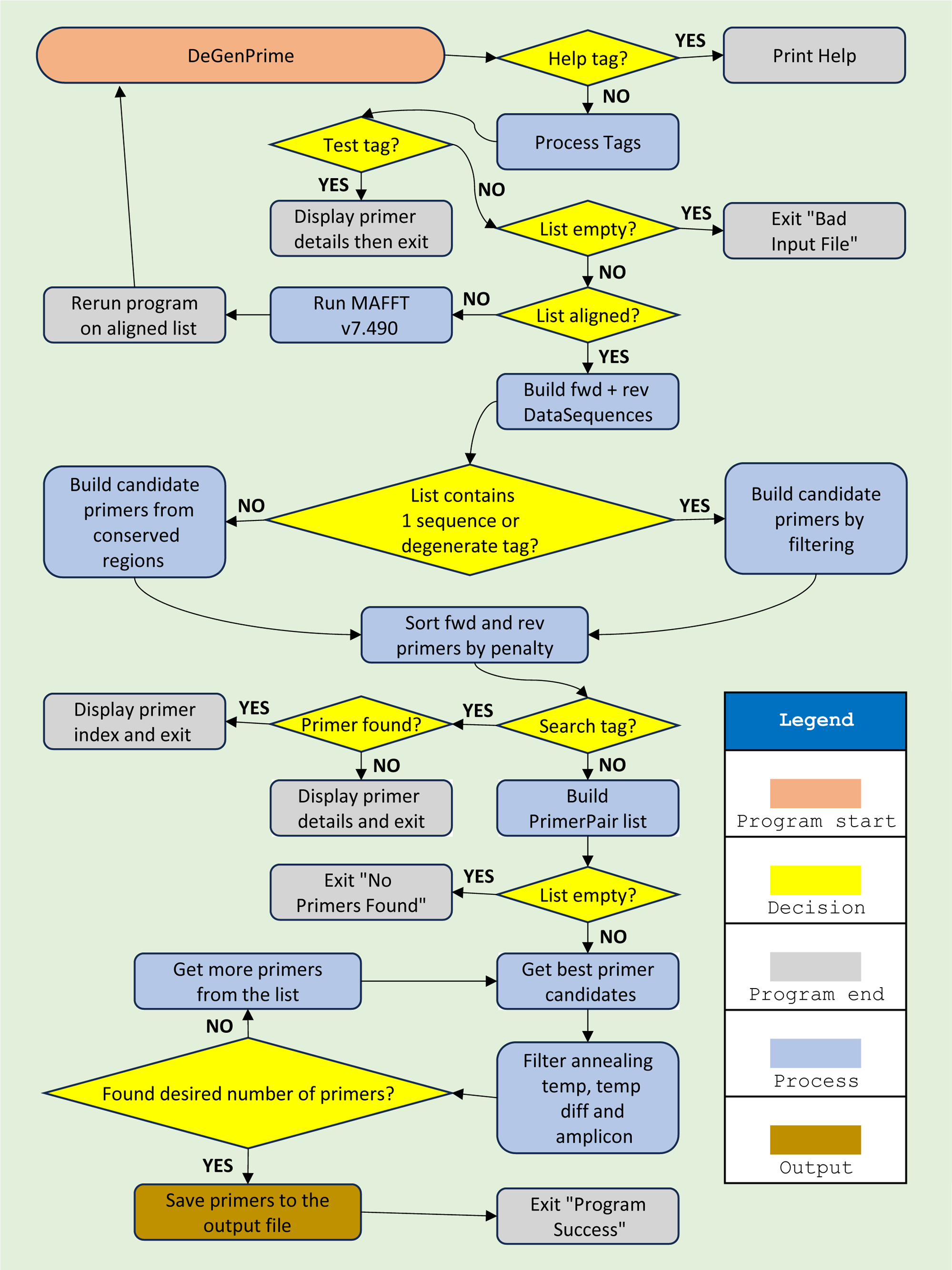
Flowgram of the DeGenPrime software. It can utilize a single sequence or a MSA as input for primer design. Input formats include a nucleotide single fasta file or a clustal MSA format. It provides two major paths for primer selection including a filtration vs. conserved region approach for primer design. DeGenPrime minimizes degeneracy but higher degeneracy can be requested upon user parameters. Outputs provided include a list of primers in .dgp format, scoring metrics, filtration results, and consensus sequence.

After these basic formatting checks are complete, DeGenPrime restructures the list of sequences into two lists of nodes representing the forward and reverse DNA sequences. Each node in this new list represents what is happening at each individual base pair (bp) location across all the sequences. It selects the most common nucleotide letter from the bp location, finds the ratio of this letter out of all possible nucleotides (A,T,G,C), and based on that ratio determines if a degenerate nucleotide code should be chosen. After this the program begins scanning the restructured list for regions where no degenerate nucleotides were identified. Regions large enough to accommodate a primer are considered conserved regions. This is our conserved region approach for MSA files by default (**Fig 1**). Once all of the conserved regions have been identified, the program runs a quick check to make sure that there are enough conserved regions or a single large conserved region able to produce a large enough amplicon (based on user specifications) for forward and reverse primers. If there are insufficient regions to find conserved primers, then the program will show the user the consensus sequence and abort. Otherwise, the possible primers within the conserved regions will be calculated and scored by primer calculator objects.

For single sequences or if the user wants some limited degeneracy, DeGenPrime can use a filtering process to generate primers instead of the conserved region approach (**Fig 1**). If a user specifies a single sequence only the filter module will be used exclusively. Forward and reverse primer calculator objects are constructed which contain lists of all possible forward and reverse primers based on the user defined parameters of if they want a minimum amplicon size or a specified range of base pairs (measuring bp 0 from the 5’ end onward). The primer calculators have built in filters for the primer lists that limit their degeneracy, deletions, GC ratio, internal repetition, melting temperature and complementary ends. Lists are filtered for primers until only the best primers remain.

The degeneracy filter is a novelty of this program. It measures the degeneracy of a single individual bp location providing the best possible base whether degenerate or not. Degenerate codes within the first or last three nucleotides within a primer will disqualify it. If any ‘N’ base is found, it is labeled as too degenerate which disqualifies a primer because it’s a 4-fold degenerate base (A,T,C,G all possible). We further limit a primer to having only one 3-fold degenerate base (e.g., H, B, V, D), and up to two 2-fold degenerate bases (e.g., S, R, Y, M, K, W) per primer. Deletions are especially problematic in primer selection and design. We designed a deletion filter that removes primers based on these rules: 1) if deletions occur within the first or last three nucleotides, 2) there are more than three consecutive deletions, 3) more than six total deletions are found, and/or 4) when the deletion causes the primer to drop below the minimum size threshold (e.g., <18 bp).

GC content must be accounted for in robust primer design. Having on average 40-60% composition per primer is required for proper Tm for the primer set. Also, having no more than 3 G or C nucleotides at the 3’ end of the primer to promote binding at annealing steps but to also limit dimerization. Our GC content filter restricts all primers regarding these parameters.

Repetitions in primers are challenging including reducing sensitivity and specificity [8]. Amplification of repetitive DNA can increase chimeras and artifacts [8]. We included a primer repetition filter based on *k-*mer counting and matching for length *K* = 2, 3, or 4 nucleotide matches that exist within the primer itself. Primers with higher matches of the variable length *K* within the filter will be excluded due to the likelihood of primer misbinding. The default for DeGenPrime based on our k-mer filter will allow for two dinucleotides, a single trinucleotide, and no tetranucleotide matches. For program efficiency, DeGenPrime does not check *k-*mers larger than four nucleotides long. Complementary ends enhance dimer formation during PCR; these must be discarded. We use a 3-mer based filter that checks the last three nucleotides of a primer for complementary ends. If a complementary match occurs within the primer, it is disqualified.

Hairpins and dimers penalty score is calculated for all sequences that have the greatest likeliness to form triloop and tetraloop hairpins [9]. Triloop hairpins occur often in a 5-bp track, where the first and last bp are complementary, and when the second bp position and the second to last bp position are G and A respectively [10]. Tetraloop hairpin follows a similar pattern as the triloop but on a 6-bp track over a 5-bp track that is complementary. Dimerization scoring takes the last 5-mer of the primer then measures the Gibbs free energy [11]. The nearest-neighbor formula for Gibbs free energy is:

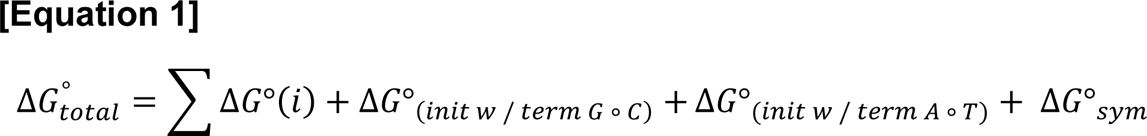

See reference [12], for equation details.

While these hairpins and dimers are not excluded from DeGenPrime they are ranked lower via penalty scoring. We use a lambda function to resolve the sorting based on the combined hairpin/dimer filter which adds a penalty to all primers with Gibbs free energy less than −3 kcal/mol or if internal nucleotides match either of these patterns.

DeGenPrime does thermodynamic calculations based on our current defaults for temperature, ion concentration, and primer concentration. All primers by default range from 50-65 °C melt temperature, 50 mM monovalent ion concentration, and 50 nM concentration of the primer itself. However, the user can specify a narrower temperature range or a selected range within our global settings. It then uses thermodynamic calculations based on the nearest-neighbor formula to determine the melting temperature of the primer. The formula is given by:

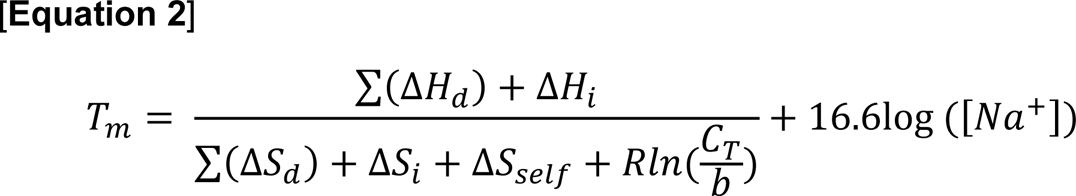

See reference [13], for equation details. Penalty is added to any primer whose melting temperature falls below the minimum or above the maximum temperature.

Within DeGenPrime, we first apply filters and scoring metrics to primer list sets (i.e., forward and reverse) independently of the primer pairing. After independent filtering and scoring of forward and reverse primers, we produce sorted lists of primers with the minimum accrued penalty first. The program then checks to see if primers were present; if not this provides an error message of ‘no primer found,’ to the user. After which we apply filtering/scoring to primer pairings.

Building a list of primer pairings and several of the filtering operations on these massive forward and reverse primer lists results in slow operations and an O(n^2^) problem. To make the program run more efficiently, we created a mapping algorithm for primer pairing (**Fig 2**). This algorithm partitions the data to ∼1,600 pairs per block, then examines them one block at a time, then moves to the next block searching for optimal primer pairs exhaustively. The program loops through until it finds five highly optimized primer pairs or results in none found. While the block size (i.e., 1,600) is arbitrary; it empirically provides a reasonable transition speed at runtime. We filter the partitions via a minimum amplicon (i.e., 100 bp default), a Tm with 1 °C between primer pairs, and annealing temperature between the pairs that are <5 °C. We calculate annealing temperature of the primer pair using this formula:

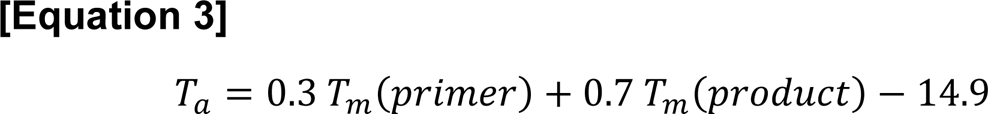

See reference [14], for equation details. If no primers are found, the user is notified. Otherwise, the final list of primer pairs is outputted to the user.

**Figure 2:**
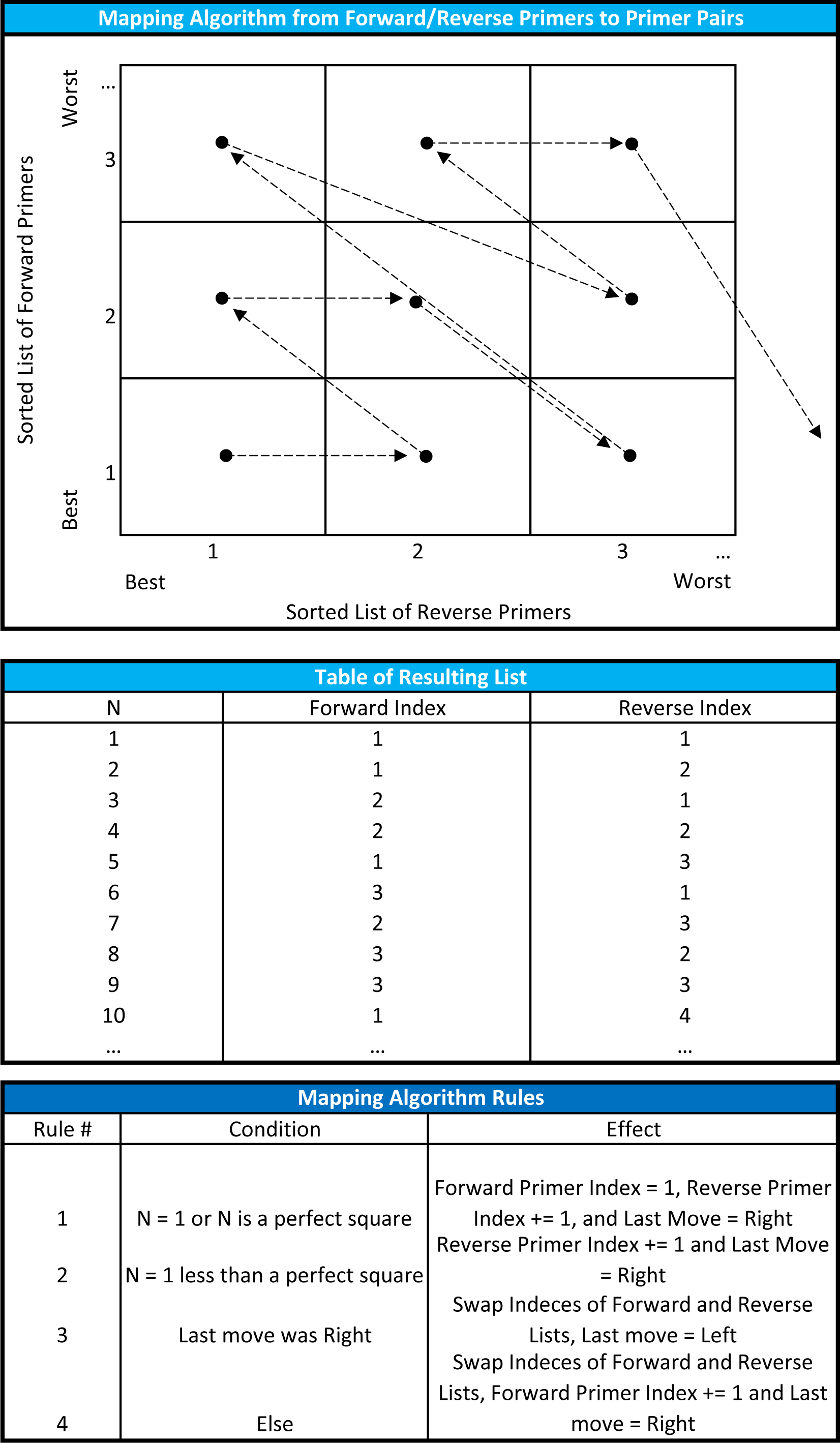
Mapping algorithm diagram. DeGenPrime utilizes a greedy loop algorithm that rank choices via step-wise mapping applying a systematic quality-control and efficient sorting of ranked lists for primer-pair optimization.

As a generalizable feature of DeGenPrime, we offer a primer testing module, to test previously assembled and designed primers or their pairs. The user can directly enter a candidate primer, then DeGenPrime performs calculations based on the aforementioned nearest neighbor formula (Equation 2), and lists whether the primer will pass various filters.

Another feature of DeGenPrime is the primer searching module. This module allows users to search for specific forward and reverse primers within their respective candidate primer lists. The program will scan each respective list for the candidate primer and alert the user if that primer was found or not found. If the primer was not found, the candidate primer is pipelined into the testing module, giving the user a clear picture of why this primer failed. Note the primer is not guaranteed to match any region within the MSA or sequence.

## 3. Results and Discussion

The novel nature of DeGenPrime makes it difficult to compare to other software. Primer3 is the top primer tool available, but its results cannot be used as a fair basis of comparison. Primer3 is only designed to process individual sequences and cannot be applied with MSA formats or support for degenerate bases like DeGenPrime. We compared Primer3 to DeGenPrime directly, finding a comparable runtime, similar penalty scoring metrics, and similar primers across both tools when utilizing one gene sequence.

EcoFunPrimer and its metagenomic version MetaFunPrimer offers an MSA processing functionality [15]. Currently, EcoFunPrimer and MetaFunPrimer cannot install, has run time errors, doesn’t compile, and currently is no longer supported. HYDEN is another and one of the first degenerate primer design softwares; however, the last update was in 2008, and it is only available in a depreciated Windows XP [16].

Some approaches to MSA primer design, like the JCVI primer designer, often make use of reference sequences [17, 18]. One problem with this approach is reference sequences introduce a primer matching bias within interspecies comparisons favoring sequences with the most matches to the given reference [19]. Within the human microbiome, where intraspecies genetic diversity and mutation rates may be higher, the reference sequence may not remain a valid basis of comparison for any particular experiment [20]. The JCVI program is no longer in active development, with the last update in 2013, and DeGenPrime is independent of a reference sequence.

We tested our program against PrimerDesign-M, which often makes use of reference sequences [21]. Currently, PrimerDesign-M is only available by web browser, not a standalone software. We utilized our primer testing module on the output from PrimerDesign-M, for iron nitrogenase (*anfD*) and found 100% of the primers failed our filters, due to including too much degeneracy (**Table 1**), and 90% failed due to having too much GC Content which shows this approach also does not consider basic primer quality filtering.

**Table 1:**
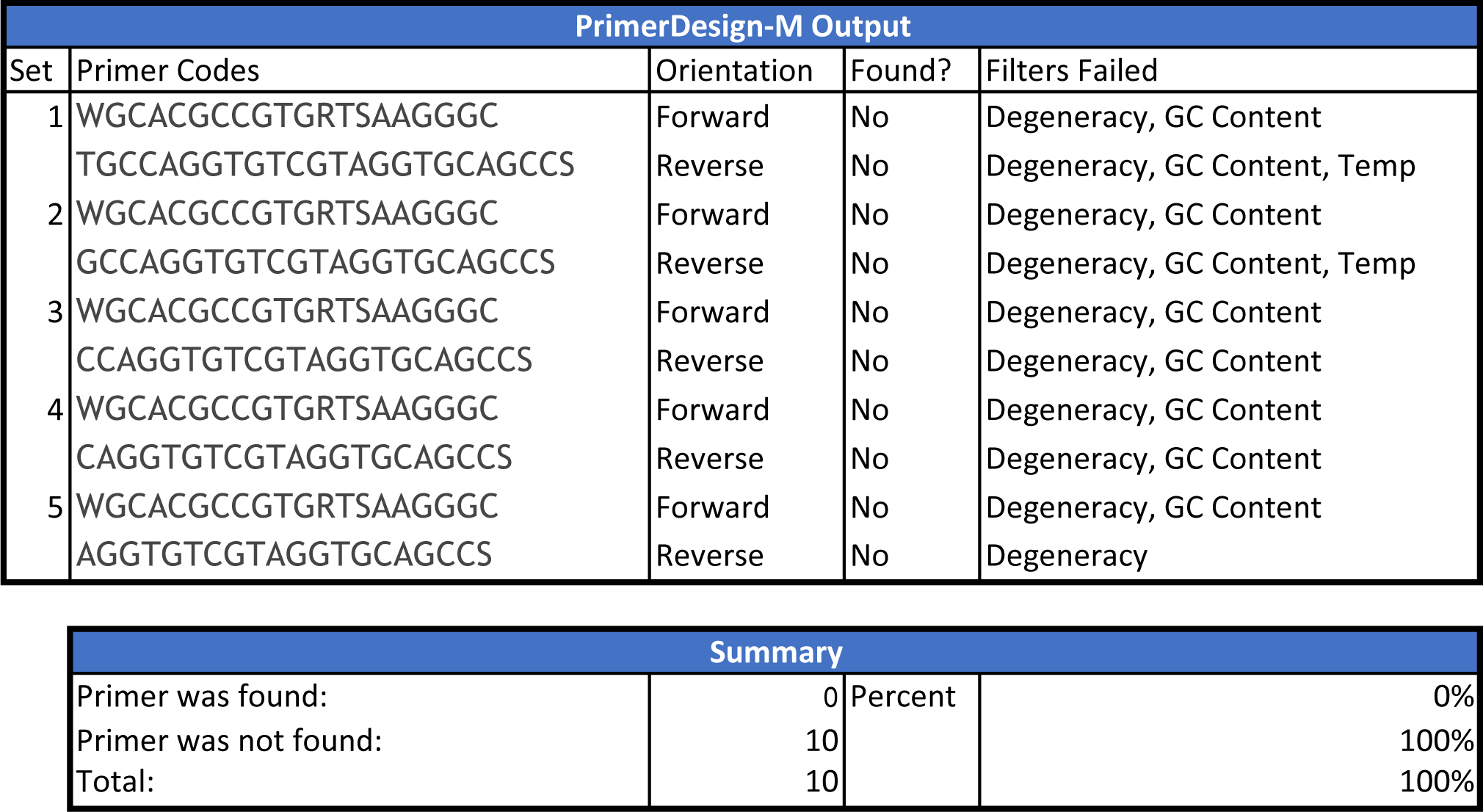
PrimerDesign-M target yield for alternative nitrogenase (*anfD*). PrimerDesign-M was ran using default parameters.

The similarly named perl program called Degeprime, provided another comparison to DeGenPrime [22]. Directly compared both programs using nitrogenase and T4-like major capsid protein (*g*23) MSA as input. We applied a search function for this direct comparison analyzing the top primers between Degeprime as a sorted list of primers. First Degeprime doesn’t report primer pairs, only forward primers making comparisons challenging. However, we scored the forward primers provided by Degeprime, ∼11% of the forward primers passed (**Table 2**), but no reverse primers, thus not a generalizable tool for robust primer design. DegePrime does not include any filtering for melting temperature ranges, GC content or complementary ends; all of which are basic criteria for making a good primer. Furthermore, some of the primers it suggested which are theoretically good primers were not located within the conserved region of the MSA. DegePrime seems to be just printing a list of all possibilities of primers without considering any quality standards.

**Table 2:**
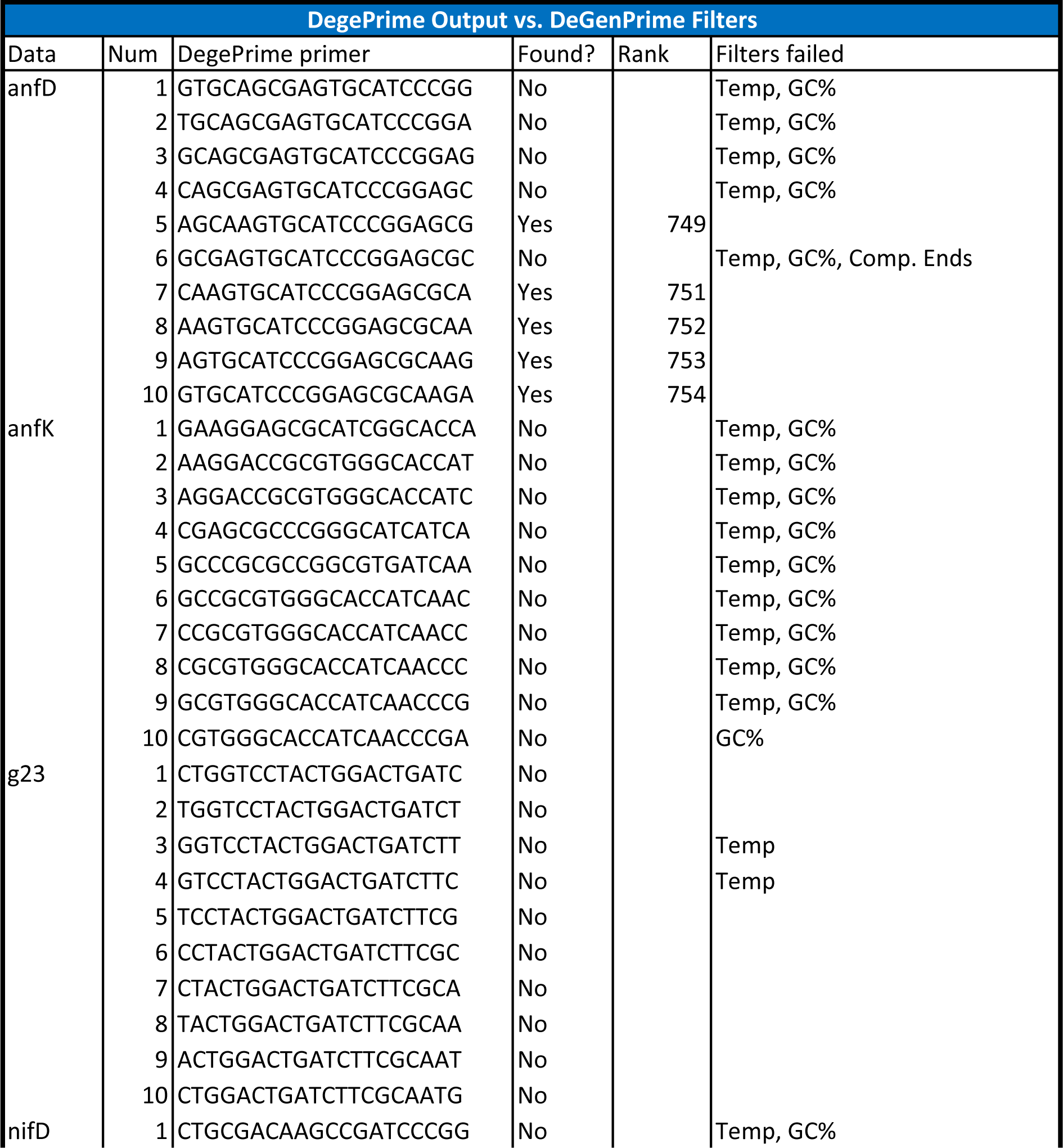
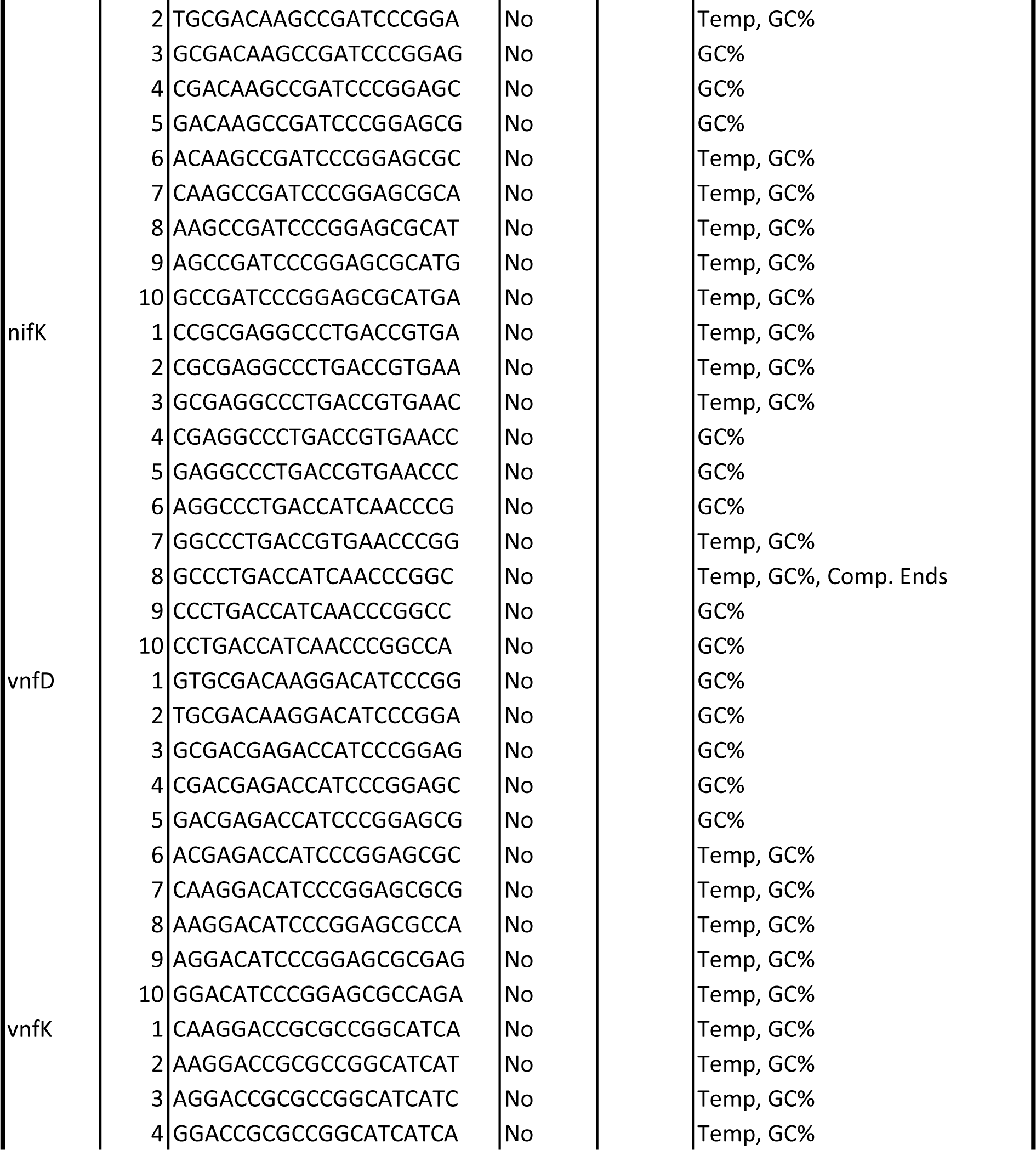
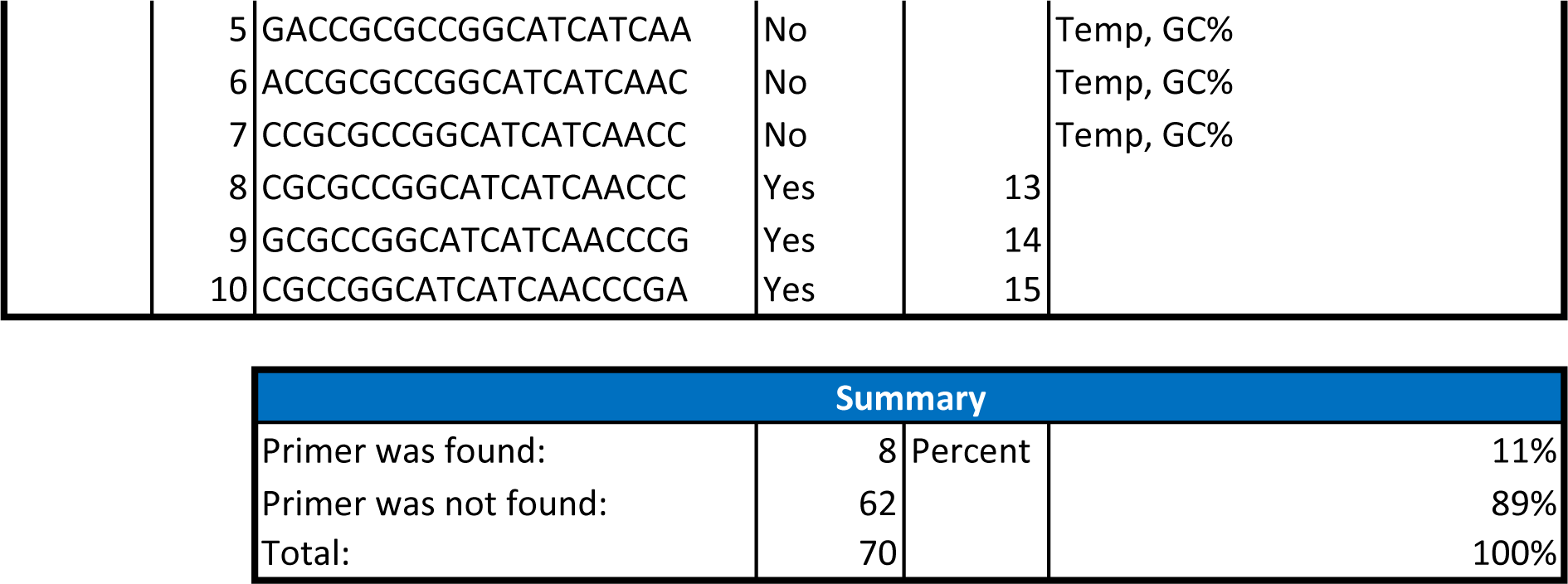
DegePrime target yield for phage major capsid (*g23*), nitrogenase (*nifDK*), and alternative nitrogenases (*anfDK/vnfDK*). DegePrime was ran using default parameters.

ConsensusPrimer is a program which, like ours, finds the conserved regions of its MSA to build its candidate list of primers [23]. It calls MAFFT to make its alignment and runs Primer3 on this alignment to find its primers. We ran each of our test MSAs into its pipeline and found subtle differences in this approach and ours.

ConsensusPrimer was unable to suggest any primers for iron-based nitrogenase (*anfK*), T4-like major capsid protein (*g23*), molybdenum-based nitrogenase (*nifD*), and vanadium-based nitrogenase (*vnfK*) alignments. The primers it did suggest for iron-based (*anfD*) and molybdenum-based (*nifK*) were slightly outside the acceptable range of DeGenPrime’s temperature filter (**Table 3**). For vanadium-based (*vnfD*), all of the reverse primers were 17 bp long which is less than the size minimum for DeGenPrime, but all of the forward primers passed filter checks. In total, 17% of primers found through this pipeline were acceptable via DeGenPrime standards, however, many of the primer pairings suggested had a forward or reverse primer repeatedly showing up in other pairings, while DeGenPrime only outputs unique primers.

**Table 3:**
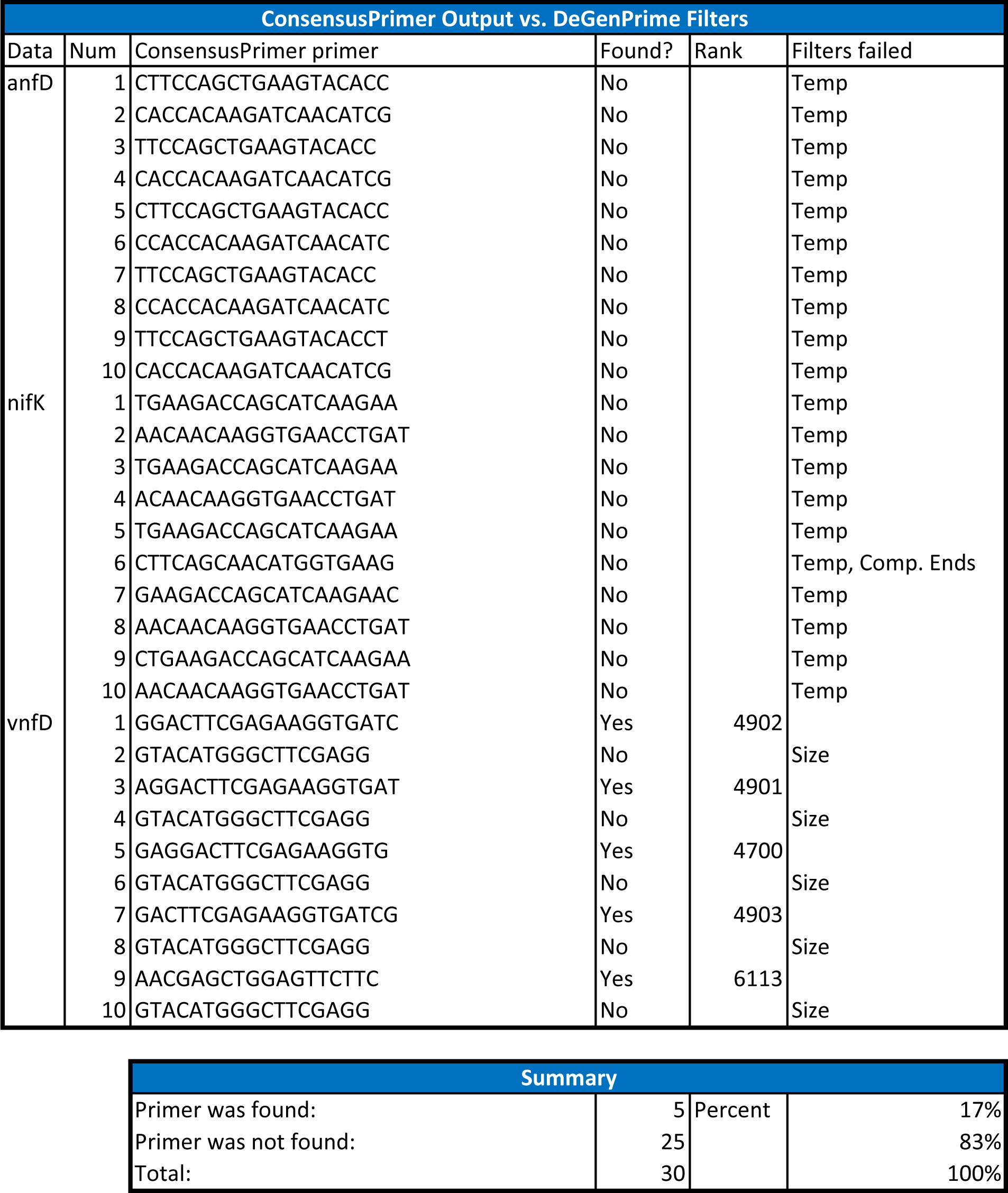
ConsensusPrimer yield for phage major capsid (*g23*), nitrogenase (*nifDK*), and alternative nitrogenases (*anfDK/vnfDK*). ConsensusPrimer was ran using default parameters. Genes *anfK, g23, nifD,* and *vnfK* yielded no primers.

A last testing method was applied to this program to see if it could produce primer pairs used in another experiment. One such experiment found degenerate primers for the T4-like major capsid protein (*g23*) phages [24]. Using the public data provided from this experiment, DeGenPrime produced forward and reverse primers which overlapped the primers used in this experiment (**Table 4**).

**Table 4:**
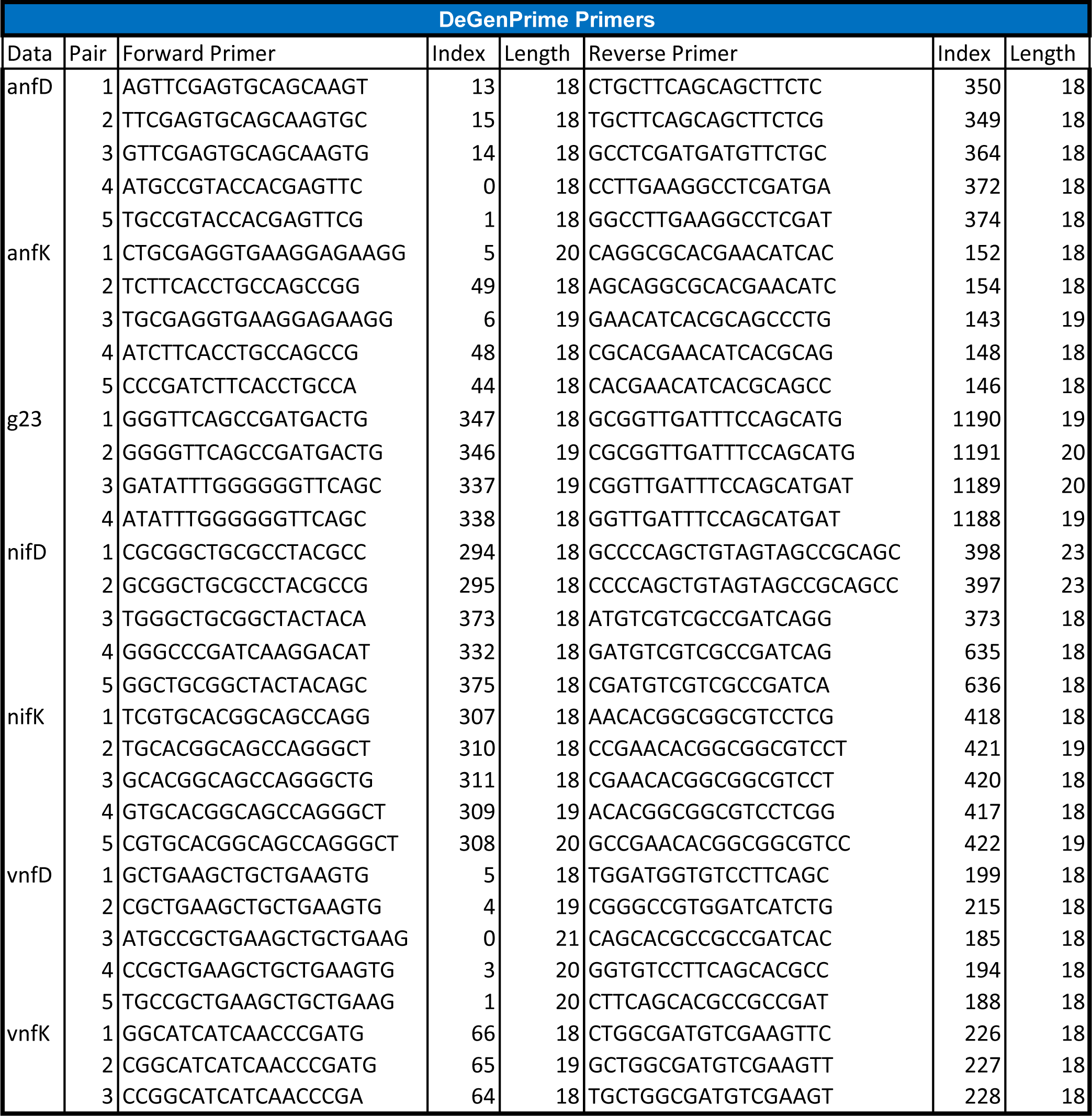
DeGenPrime yield for phage major capsid (*g23*), nitrogenase (*nifDK*), and alternative nitrogenases (*anfDK/vnfDK*). DeGenPrime was run using the conserved region approach using default.

## 4. Conclusion

The importance of having robust and accurate primer design provides time saving and financial ease for molecular diagnostics. DeGenPrime provides a novel, robust, effective approach while concurrently providing a primer evaluator. Due to the novel design of DeGenPrime it can expand the gene target amplification range of the primers which provides more targets per PCR primer set. We elucidate primer pairs while avoiding a majority of the degenerate base pairing issues. Our filtering and conversed region approaches allow for rapid primer discovery for a variety of fields within biology. Future versions will include a protein module for direct primer pair finding for aligned protein MSAs or single protein sequences. Both a GUI module and web-based versions are also currently in development providing greater accessibility to the community at large. DeGenPrime illuminates primer design unlocking the dark matter within the tree of life.

## Funding

B. W. Fulghum, S. Tanker, and R. A. White III are supported by a UNC Charlotte Bioinformatics and Genomics start-up package from the North Carolina Research Campus in Kannapolis, NC and the Department of Bioinformatics and Genomics in Charlotte, NC.

## Data Availability Statement

Raw files, code. supplemental data, and source codes, are all available on www.github.com/raw-lab/DeGenPrime.

## Acknowledgments

We acknowledge the support of the following units of the University of North Carolina at Charlotte: the College of Computing and Informatics, the Bioinformatics Research Center, the Department of Bioinformatics and Genomics, Research and Economic Development, Academic Affairs, University Research Computing and North Carolina Research Campus. We would like to thank Jessica White of the University of North Carolina Chemistry department for her assistance with the oligonucleotide thermodynamics.

## Conflicts of Interest

The authors declare no conflicts of interest. RAW is the CEO of RAW Molecular Systems (RAW), LLC, but no financial, IP, or others from RAW LLC were used or contributed to the study.

## Notes

### Competing Interest Statement

The authors have declared no competing interest.

